# Single-cell transcriptome profiling of the myeloid cells repopulating after chemotherapy identifies a neutrophil-like monocyte subset with pro-tumor activities

**DOI:** 10.1101/2024.12.28.630613

**Authors:** Zhi-chun Ding, George I. Zhou, Ogacheko D. Okoko, Xin Wang, Locke J. Bryan, Gang Zhou, Huidong Shi

## Abstract

Patients with cancer often receive chemotherapy to control tumor progression and reduce disease symptoms. Cytotoxic chemotherapeutic agents not only kill rapidly growing cancer cells but also reduce normal cells including myeloid cells, the main innate immune population involved in fighting infections and repairing tissue damages. Rapid loss of myeloid cells caused by chemotherapy triggers myelopoiesis, a process in which the hematopoietic stem and progenitor cells in the bone marrow regenerate myeloid cells, including monocytes, neutrophils, dendritic cells and macrophages, to reconstitute the myeloid cell compartment. We previously reported that chemotherapy with an alkylating agent cyclophosphamide (CTX) in mice leads to repopulation of myeloid cells that acquire immunosuppressive activities within the monocyte subset. However, detailed information on the cellular composition and molecular identity of these chemotherapy-induced immunosuppressive monocytes is lacking. Here, we investigated how the various myeloid cell subsets in the bone marrow of mice respond to CTX chemotherapy through single-cell RNA sequencing analysis (scRNAseq). We found that myeloid progenitor cells and monocytes were reduced 2 days after chemotherapy but rebounded and surpassed their pretreatment levels by day 7. Further scRNAseq analysis of pre-enriched monocytes revealed that the monocyte population was heterogenous, and that chemotherapy tilted myelopoiesis towards the production of neutrophil-like monocytes (NeuMo). We identified Cxcr4 and Cx3cr1 as suitable markers for isolation of chemotherapy-induced NeuMo and demonstrated that these cells were suppressive to T cells. Together with the evidence that CTX-induced monocytes can promote breast cancer metastasis in mice, our data reveal the heterogeneity of the monocytes reemerging after chemotherapy and identify the NeuMo subset as a potential therapeutic target for enhancing the efficacy of chemotherapy in cancer.

## Introduction

Chemotherapy is a main treatment option for patients with cancer. Chemotherapeutic agents administered to patients can kill rapidly proliferating cancer cells, leading to tumor remission. However, chemotherapy often encounters the challenge of tumor recurrence when cancer cells become resistant to further treatments. There is increasing recognition that the reactive responses of cancer and noncancer cells to chemotherapy may counteract the efficacy of chemotherapy(1). It is well known that chemotherapy can select for cancer stem cells that are inherently more chemoresistant(2). Multiple lines of evidence indicate that chemotherapy-induced changes in the composition and function of noncancer cells in the tumor microenvironment, including immune cells, endothelial cells and fibroblasts, may support the survival, metastasis, and immune evasion of tumor cells(3, 4). A better understanding of how chemotherapy alters the properties of noncancer cells is essential for the development of more effective chemotherapy regimens to achieve durable therapeutic outcomes.

Myeloid cells are a major component of the innate immune system functioning as the first line of defense against various types of pathogens. The rapid loss of myeloid cells following chemotherapy triggers myelopoiesis, a process in which the hematopoietic stem and progenitor cells (HSPCs) in the bone marrow (BM) produce new myeloid cells to replenish those lost during chemotherapy(5, 6). Myelopoiesis is a regulated multi-step process where HSPCs differentiate and develop into mature myeloid cells, including neutrophils (Neu), basophils, eosinophils, monocytes (Mo), dendritic cells (DC) and macrophages. During this process, HSPCs give rise to common myeloid progenitors (CMPs), which subsequently differentiate into granulocyte-macrophage progenitors (GMPs) and macrophage-dendritic cell progenitors (MDPs). GMPs further give rise to committed granulocytic precursors (GPs) which eventually develop into neutrophils, basophils and eosinophils. In parallel, MDPs eventually give rise to monocytes, dendritic cells and macrophages. In recent years, advances in single cell RNA sequencing technologies (scRNAseq) have revealed unprecedented insights into myeloid cell ontogeny and heterogeneity during myelopoiesis(7–9). Several studies demonstrated in mice that monocytes can derive independently from both GMPs and MDPs. In particular, the GMP-derived monocytes, which are termed neutrophil-like monocytes (NeuMo), are defined by a gene signature featuring neutrophil granule molecules, such as elastase (Elane), myeloperoxidase (Mpo), proteinase 3 (Prtn3), cathepsin G (Ctsg) and chitinase-like protein 3 (Chil3). The MDP-derived monocytes, which are termed dendritic-like monocytes (DCMo), exhibit signature genes related to MHC-II expression and DC markers. Besides differences in transcriptomic profiles, NeuMo and DCMo cells also exert distinct functions in various biological settings. NeuMo with immunoregulatory functions are found to be preferentially expanded in mice exposed to systemic lipopolysaccharide (LPS)(10, 11), and in mice subjected to brain injury(12). DCMo are found to increase in aged mice(13), and in mice after exposure to unmethylated CpG DNA(10).

Currently little is known about the relative abundance and biological function of NeuMo and DCMo in the myeloid cell compartment after chemotherapy. We previously reported that monocytes that repopulate in mice after cyclophosphamide (CTX) chemotherapy acquire a neutrophil precursor gene signature and immunosuppressive activity(14). However, whether these chemotherapy-induced immunosuppressive monocytes are equivalent to the reported NeuMo cells remains elusive. In the current study, scRNAseq analyses were conducted for whole bone marrow cells as well as highly enriched monocytes collected from mice before and after CTX chemotherapy. Our bioinformatics analysis of the scRNAseq datasets revealed the dynamic changes in various myeloid cell subsets after chemotherapy and identified a monocyte subset exhibiting the NeuMo gene signature. Our analysis also resulted in the identification of suitable markers to isolate live NueMo cells for downstream functional studies. Our data indicate that chemotherapy induces the expansion of immunosuppressive NeuMo cells that can promote tumor metastasis. These findings implicate the potential of improving the antitumor efficacy of chemotherapy by targeting the pro-tumor monocytes arising after chemotherapy.

## Methods

### Antibodies, chemicals and cell line

The following fluorochrome-conjugated antibodies, including CD11b, Ly6C, Cxcr4 and CX3CR1 were purchased from Biolegend. CellTrace Violet cell proliferation kit were purchased from Thermo Fisher Scientific. Cyclophosphamide monohydrate (CTX) was purchased from Tokyo Chemical Industry. The murine breast cancer cell line EMT6 was obtained from American Type Culture Collection (ATCC). EMT6 cells were engineered to express firefly luciferase (EMT6.luci) so that tumor cells can be detected by bioluminescence imaging (BLI). EMT6.luci tumor cells were cultured in RPMI 1640 (HyClone Laboratories) supplemented with 10% fetal bovine serum albumin (FBS), 1% penicillin/streptomycin (HyClone Laboratories), 1% non-essential amino acids and 1% glutamine (Corning) at 37 °C in a 5% CO2 incubator.

### Bone marrow cell preparation for flow cytometry analysis

Tibia and femur bones were collected from euthanized mice. Bone marrows were collected by flushing the bones with PBS. After lysing red blood cells by Ammonium-Chloride-Potassium (ACK) buffer, bone marrow single-cell suspensions were stained with fluorochrome-conjugated antibodies for 15 minutes at room temperature in the dark. All flow cytometry analysis data were acquired on an Attune NxT Flow Cytometer (Invitrogen) and analyzed using Flowjo software (Tree Star).

### In Vitro T cell suppression assay

Spleens collected from normal BALB/c mice were grinded thoroughly in PBS and the suspensions were passed through a 70 μM nylon mesh to remove debris. After lysing red blood cells with ACK buffer, spleen cells were labeled with 0.1 μM CellTrace violet dye and seeded in triplicates into a round-bottom 96-well plate with 1 × 10^5^ cells per well in 200 µl medium. T cells (both CD4+ and CD8+) within spleen cells were stimulated with 1 µg/ml of anti-CD3 (Biolegend, clone 145-2C11) and 5 µg/ml of anti-CD28 (Biolegend, clone 37.51). 1 × 10^5^ NeuMo cells or other classical monocytes (cMo) sorted by fluorescence-activated cell sorting (FACS) were added to specified wells. Cells were harvested 3 days after culture for flow cytometry analysis.

### Mice

Balb/c mice of 4 to 6 weeks of age were purchased from Charles River Laboratories. All mice were housed under specific pathogen-free conditions by Laboratory Animal Services of Augusta University. All animal experiments were approved by the Institutional Animal Care and Use Committee (IACUC) of Augusta University.

### Administration of chemotherapy, monocytes infusion and tumor inoculation to mice

For chemotherapy with cyclophosphamide, CTX powder was dissolved in PBS and intraperitoneally injected to mice at the dose of 150 mg/kg. At the specified time, mice were euthanized to collect bone marrow cells from the tibia and femur bones. Bone marrow cells were removed of red blood cells using the Ammonium-Chloride-Potassium (ACK) buffer. Bone marrow cells were stained with antibodies against CD11b and Ly6C and subjected to cell sorting using the Invitrogen Bigfoot Spectral Cell Sorter. The sorted monocytes (CD11b^+^Ly6C^hi^) or neutrophils (CD11b^+^Ly6C^intermediate^) were intravenously injected to recipient mice through tail vein. Each mouse received 0.8 – 1.0 million sorted monocytes or neutrophils weekly for a total of four infusions, with the first infusion performed right before tumor inoculation. For tumor inoculation, EMT6.luci tumor cells were harvested and washed with phosphate-buffered saline (PBS). Tumor cells were inoculated to the 4th mammary fat pads of female BLAB/c mice with each mouse receiving 0.1 million EMT6.luci cells in 50ul volume. The tumor size was measured by caliper every other day and expressed as the product of two perpendicular diameters in square millimeters.

### Bioluminescence imaging

Bioluminescent imaging (BLI) was performed on a Spectral Advanced Molecular Imaging X (Ami X) system (Spectral Instruments Imaging) to detect the presence of the primary and metastasized tumor cells. For live mice imaging, each mouse received an intraperitoneal injection of 150 mg/kg luciferin and was anesthetized by inhalation of 2% isoflurane. The photographic images were acquired and overlaid with pseudocolor luminescent images. For imaging of the metastasized tumors, mice were euthanized and immediately dissected to expose the internal organs. The primary tumors were resected and taken away from the mice to prevent interference with detection of metastases in distant organs. All BLI data were analyzed using the Aura Imaging Software (Spectral Instruments Imaging). The luminescence signal in the tumor region was quantified as photon/sec as an indicator of tumor burden.

### Cell preparations for scRNAseq

For scRNAseq of whole bone marrow cells, bone marrows were collected from cyclophosphamide-treated mice on day 2 (CTX_D2) and day 7 (CTX_D7). Bone marrows from untreated mice were used as the control (naïve). Live cells were enriched using the EasySep Dead Cell Removal kit (Stemcell Technologies). For scRNAseq of pre-enriched monocytes, bone marrow cells were stained with anti-CD11b and anti-Ly6C antibodies and sorted for CD11b^+^Ly6C^hi^ monocytes by Bigfoot Spectral Cell Sorter. The purity of sorted monocytes was normally greater than 98%.

### Single-cell RNA sequencing

Freshly isolated whole bone marrow cells or monocytes (CD11b^+^Ly6C^hi^) enriched by fluorescence-activated cell sorting (FACS) were processed for scRNAseq libraries using the Chromium Controller (10X Genomics). scRNAseq libraries were generated for approximately 2,000 cells per sample using 10X 3’ single cell mRNAseq V3 reagents. The scRNAseq libraries were sequenced using Illumina NextSeq500 sequencer to collect approximately 80k reads per cell. Each cell was tagged with a 16bp barcode sequence, which represents the identity of each single cell throughput of the analysis pipeline.

### scRNAseq data analysis

The raw reads in the fastq format were processed using the 10X genomics cellranger analysis package. Samples were pooled using the cellranger aggr pipeline. The cellranger pipeline outputs containing gene-by-cell expression data were processed using the Python package scanpy. Graphs were generated using plotting functions available in the scanpy, Pandas, and Matplotlib packages. Data quality control measures were implemented using scanpy, and only cells with at least 200 genes and less than 20% of mitochondria gene content were retained for further analysis. Normalization, scaling, dimension reduction, integration, and clustering were carried out using scanpy. Dimension reduction was implemented with umap to simplify 40,000 dimensions to readable graphs, with clusters close together having similar cells. Top differentially expressed marker genes for each cluster were identified by scanpy built-in functions. The myeloid precursors and mature myeloid cells in bone marrow were identified by gene signatures published by Kwok et al(15). NeuMo and DCMo subsets in the sorted monocyte population were identified by gene signatures published by the Goodridge’s group(10, 13).

### RNA velocity analysis

RNA velocity analysis was performed using the Python package scvelo, which employs a modified dynamical model to relate the dynamics of unspliced pre-mRNA and spliced mature RNA. The bam files generated by the cellranger pipeline were pre-processed using the velocyto command- line tool (CLI) with default parameters specific for 10X genomics data. RNA velocity was estimated using default parameters in scvelo. Velocity graphs were generated using functions in scvelo and scanpy based on the UMAP embedding generated by scanpy analysis.

### Statistical analysis

Data were analyzed using Prism 7.0 (GraphPad Software, Inc.). For comparison between two groups, the statistical significance of the results was determined using unpaired two-tailed Student’s *t* test. One-way ANOVA was used to determine statistical differences among three or more groups. For multiple testing correction in sequencing analyses, the false discovery rate (FDR) method was used. The adjusted p-value described in this manuscript refers to FDR.

## Results

### scRNAseq analysis reveals the dynamic changes in myeloid cell composition during CTX-induced myelopoiesis

Since our previous work show that CTX-induced monocytes have immunosuppressive activity(14), it is important to understand how this population evolves along with other cells in the bone marrow following chemotherapy. To have a comprehensive understanding of the composition of the myeloid cell compartment during CTX-induced myelopoiesis, we conducted scRNAseq analysis for bone marrow samples collected from mice 2 and 7 days after CTX treatment, with corresponding samples from untreated naïve mice as the control (Fig. 1 schema). From a total of 22,862 cells analyzed (7,649 cells from naïve, 6,608 cells from CTX-D2, 8,605 cells from CTX_D7), we identified 21 cell clusters with each cluster annotated to a specific cell type (Fig. 1A). The gene signatures used for cell cluster annotation were listed in the heat map shown in Fig. 1B. Analysis of cell composition changes over time after chemotherapy indicates that myeloid cells, mainly monocytes (cluster 6) and myeloid progenitor cells, including GMP (cluster 8), MDP (cluster 9), progenitor neutrophils (proNeu - cluster 3) and pre-neutrophils (preNeu - cluster 4), were largely depleted 2 days after CTX, while T and B lymphocytes remained intact (Fig. 1C). By day 7 after CTX treatment, all myeloid cell subsets had returned to the pre-treatment level, with monocytes and myeloid progenitor cells (GMP, MDP, proNeu and preNeu) showing increased fractions in the bone marrow at the expense of lymphocytes such as T and B cells. The results indicate that myelopoiesis occurs prior to lymphopoiesis after CTX treatment, and that monocytes repopulate quickly (within 7 days) in the bone marrow. We next performed RNA velocity analysis, in which the transcription kinetics is estimated by distinguishing the spliced and unspliced mRNA and used to predict the future state of individual cells on the time scale of hours. As shown in Fig. 1D, GMP gave rise to neutrophils through the proNeu-preNeu- immature neutrophil pathway, while MDP gave rise to monocytes and DCs in the bone marrow. The results confirm the lineage hierarchy of the myeloid cells, validating the quality of our dataset and the reliability of my cell cluster identification and annotation.

**Figure. 1.**
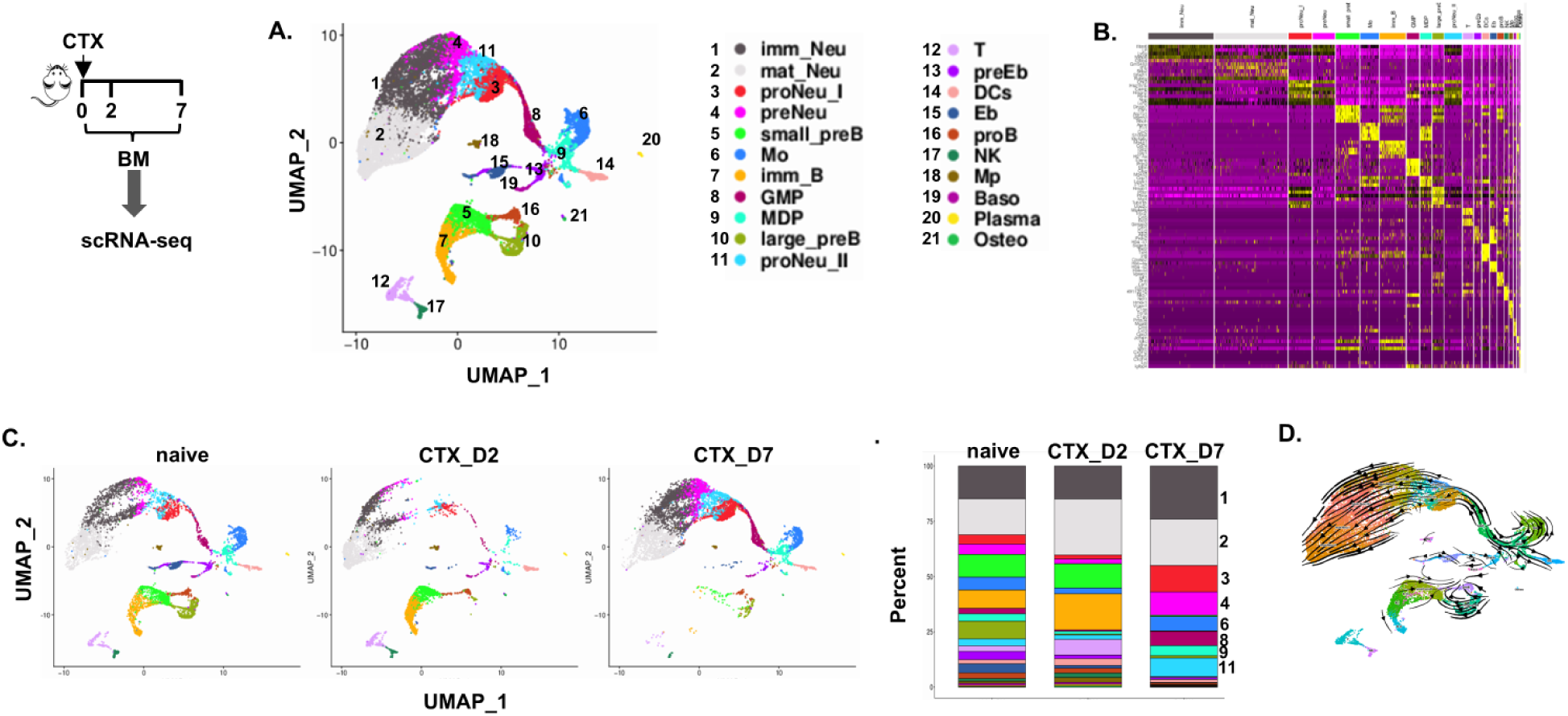
scRNAseq analysis reveals the dynamic changes in immune compositions in the bone marrow following chemotherapy with CTX. The procedures and timeline are depicted in the schema. Briefly, bone marrow (BM) cells were collected from CTX-treated mice on day 2 and day 7. Bone marrow cells from naïve mice were used as control. After removing dead cells, the whole bone marrow cells were subjected to 10x Genomics scRNAseq analysis. A. The Uniform Manifold Approximation and Projection (UMAP) visualization of 21 cell clusters that are annotated based on gene expression profiles. B. Heatmap of top 10 genes in each cluster. The columns represent cells and the rows present genes. Cells are grouped by clusters and top 10 most significant markers are shown. C. The UMAPs of each samples indicate the dynamic changes of each cluster after chemotherapy. The percentages of each cluster in the specified bone marrow samples are summarized in the bar graph. The cell clusters marked by the side of the bar graph indicate those clusters showing increased percent in CTX-D7 bone marrow compared to naïve bone marrow. D. RNA velocity analysis confirms the lineage relationship of the cell clusters.

### scRNAseq analysis of CTX-induced monocytes reveals a heterogeneous population containing cell subsets with gene signatures resembling NeuMo and DCMo respectively

Published studies have uncovered the heterogeneity of the classical monocytes (CD11b^+^Ly6C^hi^) and identified at least two subsets, i.e. NeuMo and DCMo, based on transcriptomic profiling(7, 10). Although our scRNAseq analysis of the whole bone marrow cells was highly informative in revealing the impact of chemotherapy on the myeloid cell compartment, the limitation in cell numbers prevented in-depth examination of the monocyte population. To gain higher resolution on monocyte heterogeneity, we performed scRNAseq analysis on pre-enriched CD11b^+^Ly6C^hi^ classical monocytes collected from the bone marrow of naïve mice and mice receiving CTX 7 days ago (Fig. 2 schema). A total of 4,337 cells were analyzed, with 1,846 cells for the naïve sample and 2,491 for the CTX_D7 sample. By applying the NeuMo and DCMo gene signatures reported by Barman et al(13) to our scRNAseq dataset, we identified 2 clusters of cells (cluster 2 and cluster 3) expressing high levels of NeuMo signature genes, including (Mpo, Elane, Prtn3 and Ctsg) (Fig. 2A-B). Cluster 3 NeuMo cells showed higher levels of genes involved in cell proliferation (Mki67, Hmgb2, Ptma, Plaf and Tubb5) compared to Cluster 2 NeuMo cells. I also identified a minor subset (cluster 6) with the gene signature characteristic of DCMo (CD74, H2-Aa, H2-Ab1 and Batf3), along with other classical monocytes with less defined gene signatures population with undefined gene signature. Fig. 2C shows that NeuMo and DCMo subsets exist in the bone marrow of naïve mice, but the fraction of NeuMo (the combination of cluster 2 and cluster 3) increased in the bone marrow of mice 7 days after CTX treatment. The results demonstrate for the first time that chemotherapy-induced myelopoiesis leads to the expansion of monocytes with a NeuMo identity.

**Figure. 2.**
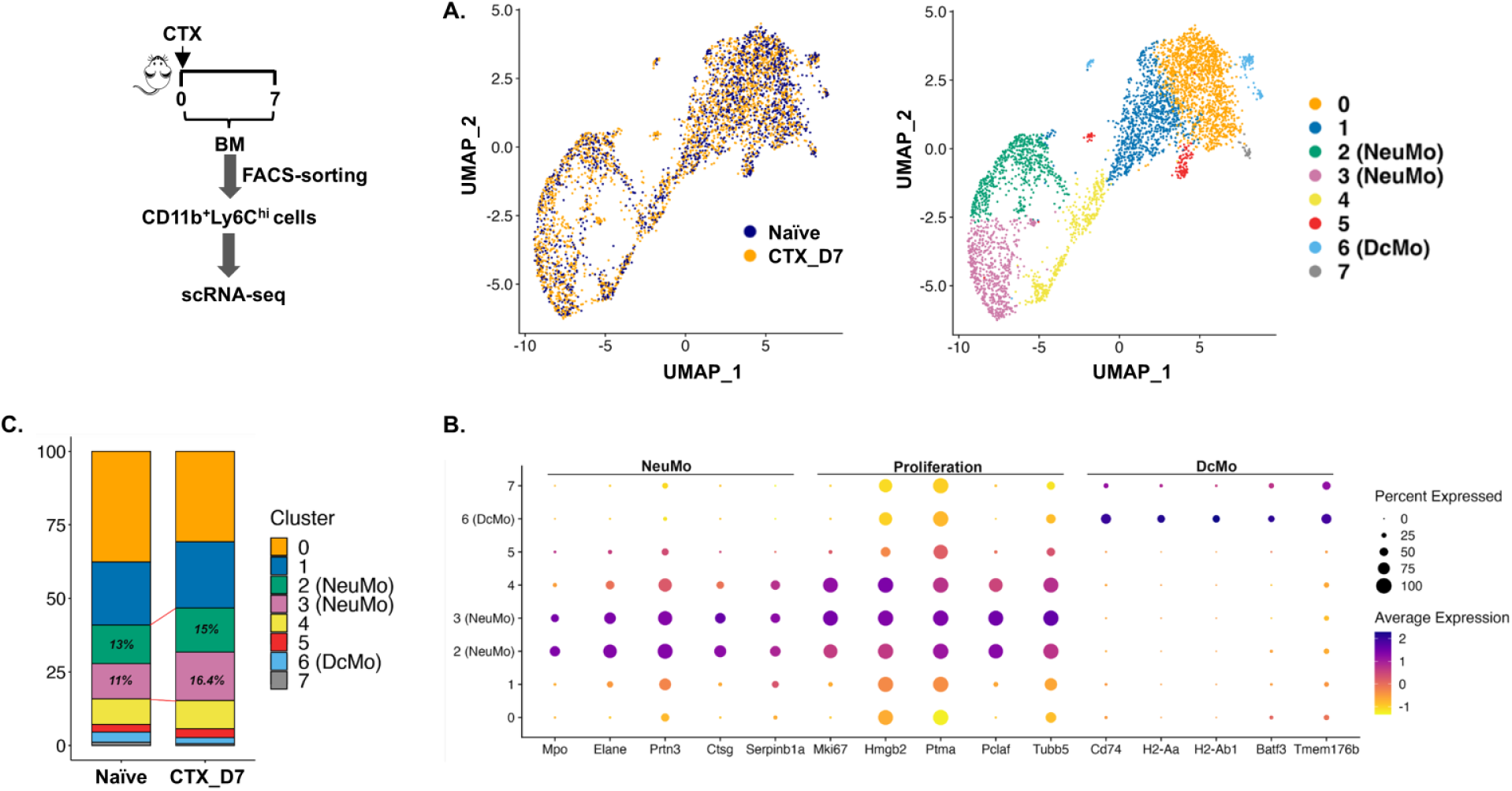
scRNAseq analysis of pre-enriched monocytes identifies distinct cell subsets based on gene expression profiles. The experimental procedures and timeline are depicted in the schema. Briefly, bone marrow (BM) cells were collected from CTX-treated mice on day 7. Bone marrow cells from naïve mice were used as control. Monocytes (CD11b^+^Ly6C^hi^) were FACS-sorted to high purity for scRNAseq analysis. A. UMAP shows distribution of cells from different samples (left panel) and cell cluster/cell subset identification (right panel). The signature genes used for cell clustering are outlined in the dot plots shown in B. C. The percentages of each cluster in the specified bone marrow samples are summarized in the bar graph.

### Identification of surface markers for the enrichment of NeuMo cells

We intended to identify suitable markers that would allow us to enrich NeuMo cells for functional studies. Based on the analysis results of our monocyte scRNAseq dataset, we selected Cxcr4 and Cx3cr1 as potential markers because they had largely nonoverlapping expression distribution in NeuMo and the rest of the classical monocyte population (Fig. 3A), and they encode surface molecules that can be used for FACS-sorting of live cells. By flow cytometry analysis, it turned out that the combination of CD11b, Ly6C, Cxcr4 and Cx3cr1 provided a reasonable resolution for separating NeuMo cells from other monocytes (Fig. 3B). The NeuMo cells (CD11b^+^Ly6C^hi^Cxcr4^+^Cx3cr^-^) appeared to have increased size (FSC) and granularity (SSC) compared to other monocytes (CD11b^+^Ly6C^hi^Cxcr4^-^Cx3cr^+^ and CD11b^+^Ly6C^hi^Cxcr4^-^Cx3cr^-^) (Fig. 3C), in line with the results of published studies(10, 13). Consistent with the scRNAseq analysis (Fig. 2), our antibody staining panel confirmed that CTX treatment tilted myeloipoiesis towards production of more NeuMo cells, as reflected by a increased ratio of NeuMo over other classical monocytes (NeuMo/cMo) within the bone marrow monocyte population after chemotherapy (Fig. 3D).

**Figure. 3.**
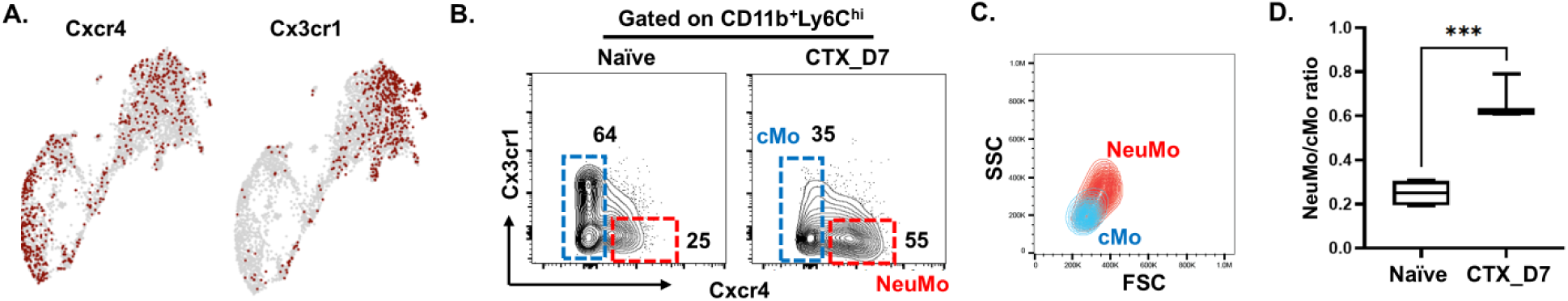
Identification of cell surface markers that distinguish CTX-induced NeuMo cells. A. UMAP distribution of the candidate genes. The candidate genes are selected because their expressions are enriched in NeuMo (Cxcr4) or the rest of the classical Ly6C^hi^ monocytes (Cx3cr1), and they encode surface molecules that can be used for live cell isolation. B. The combination of Cxcr4 and Cx3cr1 can be used to distinguish NeuMo from other classical monocytes (cMo) within bone marrow monocyte population. The numbers in dot plots demarcate the percents of NeuMo and other monocytes in bone marrow-derived monocytes. C. NeuMo cells show increased size (FSC) and granularity (SSC) compared to other monocytes. D. The ratio of NeuMo over other monocytes increases in bone marrow after CTX treatment. The ratio is calculated based on the data shown in B and shown as mean±SEM with 3 mice in each group. ***, P<0.001. FSC, forward side scatter; SSC, side scatter.

### The immunosuppressive activity of CTX-induced monocytes is mediated by the NeuMo subset

We next investigated which monocyte subset possessed the immunosuppressive activity we previously observed with the CTX-induced monocyte population. NeuMo and other classical monocytes (cMo) from the bone marrow of CTX-treated mice were FACS-sorted based on their CXCR4 and CX3CR1 expression profiles (Fig. 4A schema). The sorted monocytes were used for *in vitro* T cell suppression assays under the conditions specified in Fig. 4A. Specifically, splenocytes from naïve mice, which contained both CD4+ and CD8+ T cells, were labeled with CellTrace violet dye and used as responder cells. The responder cells were cultured alone or in co-culture with the FACS-sorted monocytes, in the presence of anti-CD3 and anti-CD28 antibodies as T cell activation stimuli. Splenocytes cultured alone without stimulation were used as a negative control. Cells were harvested for flow cytometry analysis 3 days after culture. Upon stimulation, activated T cells would undergo several rounds of cell division and upregulate CD25 expression. Therefore, the higher fraction of divided (violet-diluted) CD25^hi^ cells in the responder would indicate stronger T cell activation. Fig. 4B-C shows that the presence of CTX-induced none- NeuMo classical monocytes (cMo) did not reduce T cell activation compared to T cells stimulated alone. Notably, CTX-induced NeuMo cells significantly reduced T cell activation in both CD4+ and CD8+ T cell subsets. Interestingly, NeuMo cells isolated from the bone marrow of naïve mice did not inhibit T cell activation. These results indicate that the immunosuppressive activity of CTX-induced monocytes resides predominantly in the NueMo subset, and that NeuMo cells acquire this feature during myelopoiesis after chemotherapy.

**Figure. 4.**
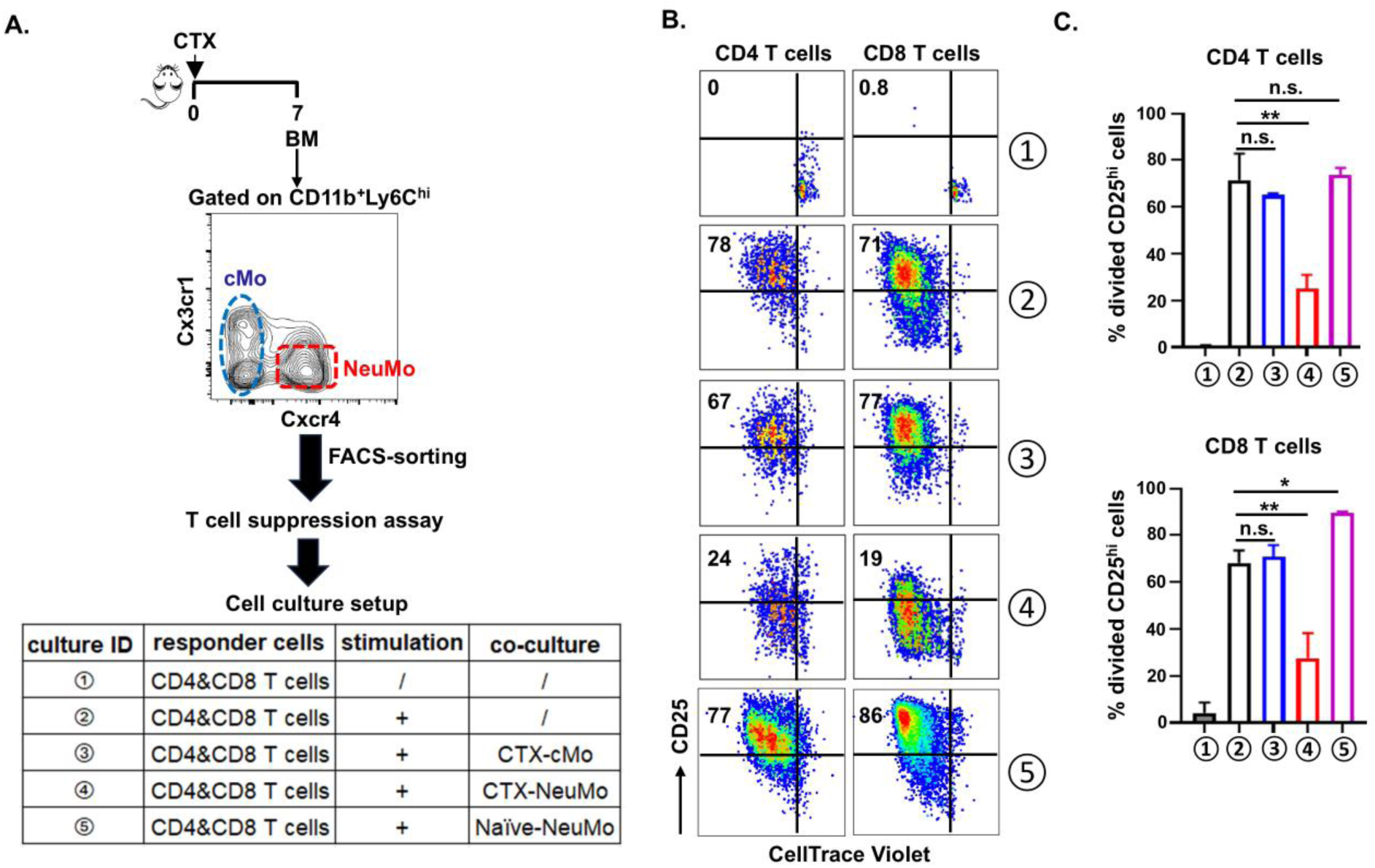
CTX-induced NeuMo cells but not DCMo cells can suppress T cell activation. A. The schema depicts the experimental procedures and timeline. 7 days after CTX treatment, bone marrow monocytes (CD11b^+^Ly6C^hi^) were subjected to FACS-sorting to isolate NeuMo (Cxcr4^+^Cx3cr1^-^) and other classical monocytes (Cxcr4^-^) cells. The sorted monocytes were used for *in vitro* T cell suppression assay in which T cell-containing splenocytes from naïve mice were labeled with CellTrace violet dye and used as responder cells. The cell culture set up is outlined in the chart. 3 days after culture, cells were harvested and stained with antibodies against CD4, CD8 and CD25. T cell division and activation were evaluated by dilution of violet dye intensity and upregulation of CD25, respectively. B. Representative dot plots are gated on CD4+ (left panel) and CD8+ (right panel) T cells to show cell division status and CD25 expression level. Numbers in dot plots represent percent of fully activated T cells (divided CD25^hi^) under the specified culture condition. The results are summarized in the bar graphs in C shown as mean±SEM of triplicate samples. *, P<0.05; **, P<0.01; n.s., non-significant.

### CTX-induced monocytes can promote tumor metastasis

It has been reported that NeuMo cells induced under LPS-induced inflammatory condition can promote tumor metastasis(11). We asked whether CTX-induced NeuMo cells have similar pro-tumor activity. To address this, we used an orthotopically implanted breast cancer model to examine whether adoptive transfer of CTX-induced monocytes can promote tumor metastasis. Due to the technical difficulty of sorting large numbers of NeuMo cells for adoptive transfer, we decided to transfer the whole monocyte population, which contained increased proportion of NeuMo cells. As shown in Fig. 5A schema, bone marrow cells were collected from mice 7 days after CTX treatment and subjected to fluorescence-activated cell sorting to isolate monocytes (CD11b^+^Ly6C^hi^), with neutrophils (CD11b^+^Ly6C^int^) sorted simultaneously for comparison purpose. These sorted myeloid cells were intravenously injected to female BALB/c mice, which were subsequently implanted with luciferase-expressing breast cancer cells EMT.6 (EMT6.luci) in the mammary fat pads. Infusion of sorted monocytes and neutrophils was repeated weekly for a total of 4 injections. Mice implanted with EMT6.luci without myeloid cell infusion were used as a control. The use of EMT6.luci cells would allow sensitive detection of tumor cells at the primary site (breast) by live mice bioluminescence imaging (BLI), as well as in distant organs (metastases) by *ex vivo* BLI at the time of animal euthanization. The growth kinetics of the primary tumors was determined by caliper measurement of the tumor area (length x width) every other day. Infusion of CTX-induced myeloid cells, either the neutrophils of monocytes, did not affect the growth of the primary tumors because the tumor burdens did not differ between groups as reflected by both live mice BLI (Fig. 5B) and tumor size measurement (Fig. 5C). At the endpoint of the experiment, metastases were not detected by *ex vivo* BLI in any distant organs in mice of the tumor only control group, as expected for EMT6 as a nonmetastatic tumor. Notably, tumor signals were detected in distant organs, including lung, liver and intestine, in mice receiving infusion of CTX-induced monocytes but not neutrophils (Fig. 5D). Taken together, the results indicate that CTX-induced monocytes, most likely the NeuMo subset, can promote tumor metastasis.

**Figure. 5.**
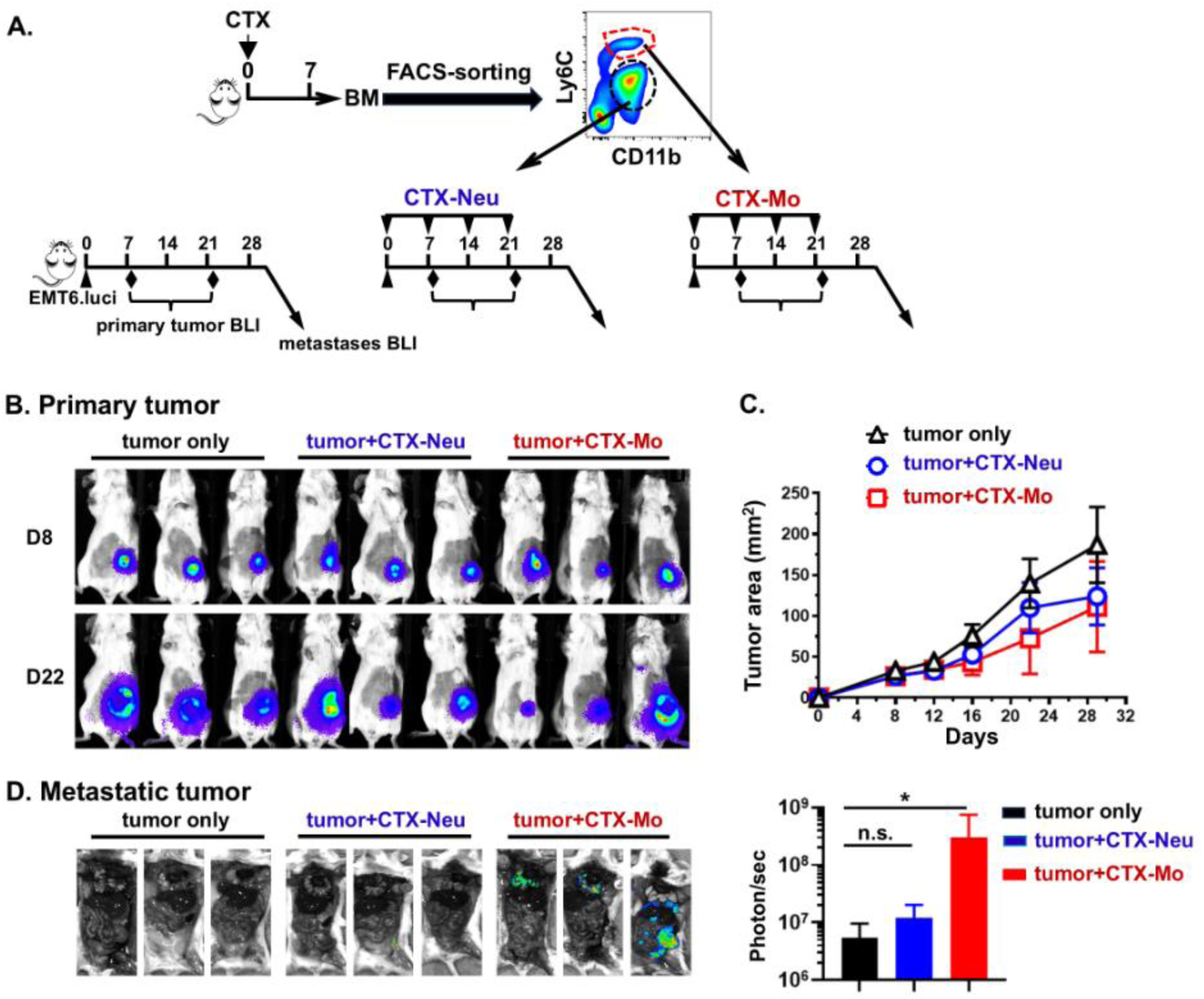
CTX-induced monocytes but not neutrophils promote tumor metastasis. A. The experimental procedures and timelines are depicted in the schema. Briefly, naïve BALB/c mice received intraperitoneal injection of cyclophosphamide (CTX) at the dose of 150 mg/kg. 7 days later, bone marrow (BM) cells were collected from these mice for staining with CD11b and Ly6C antibodies. Neutrophils (CTX-Neu) and monocytes (CTX-Mo) were FACS-sorted to high purity and infused to recipient mice via tail vein injection. Mice were implanted with EMT6.luci cells in the mammary fat pads after myeloid cell infusion. One cohort of mice received tumor inoculation only was included as control. Infusion of FACS-sorted CTX-induced monocytes or neutrophils was repeated on day 7, day 14 and day 21. Tumor sizes were measured by caliper every other day. On day 8 and day 22, the burdens of the primary tumors were measured by BLI. All mice were euthanized on day 30, dissection was performed to remove the primary tumor and expose the internal organs of each mouse for detection of metastases in distant organs by BLI. B. The burdens of the primary tumors were evaluated by BLI at the specified time points. C. Tumor areas (length x width) were plotted against time to present tumor growth curves shown as mean±SEM. The three experimental groups show overlapping tumor growth curves of the primary tumor. D. Detection of metastases in distant organs by BLI on day 30. Results of luciferase signal intensity (photon/sec) in distant organs quantified as mean±SEM are summarized in the bar graph at right. *, P<0.05; n.s., non-significant.

## Discussion

Bone marrow is a tissue where hematopoietic cells are generated and give rise to myeloid cells and lymphoid cells, cells that protect us from diseases such as infections and cancer. Chemotherapeutic agents used to kill rapidly growing cancer cells can also affect normal cells in the bone marrow, including myeloid cells. Myeloid cell regeneration, a process referred to as myelopoiesis, takes place in the bone marrow to supplement the normal cells lost to chemotherapy(5, 6).

Our single-cell RNAseq data analysis reveals the dynamic changes in cell compositions in the bone marrow after chemotherapy. We found that neutrophil precursors, monocytes and granulocyte-monocyte progenitors (GMPs) were markedly reduced shortly after chemotherapy (day 2). However, these cells rebounded afterwards and suppressed their pretreatment levels in the bone marrow by day 7. In contrast, reduction of lymphocytes, including B cells, T cells and NK cells, did not occur on day 2, but became evident by day 7 (Fig. 1C). The faster recovery of myeloid cells may reflect the attempt of the host to restore the number of cells that serve as the fist-line of defense against invading pathogens.

We previously reported that CTX chemotherapy induces monocytes with immunosuppressive activity(14). These monocytes express some markers characteristic of neutrophil precursors, including Mpo, Elane, Prtn3 and Ctsg. This feature prompted us to investigate whether these cells are equivalent to the so-called neutrophil-like monocytes (NeuMo) identified under certain infection or inflammation settings. Our scRNAseq analysis confirms the presence of NeuMo and DCMo subsets in CTX-induced monocytes based on gene expression profiles. We show that NeuMo and DCMo cells are present in mouse bone marrow in steady state. Notably, the NeuMo cells expanded after chemotherapy, not the NeuMo cells from naïve mice, possess the ability to suppress T cells (Fig. 4), indicating that immunosuppressive activity is an acquired feature. Future work should investigate the mechanisms of NeuMo’s transition from nonsuppressive to suppressive in the post-chemotherapy window.

A recent study reported that chemotherapy with gemcitabine or paclitaxel plus doxorubicin induces accumulation of monocytes in the lungs to drive metastasis(16). Interestingly, the monocytes that promote tumor metastasis in this study also express several NeuMo signature genes, such as Prtn3 and Elane. It is likely that expansion of NeuMo cells is a general feature of myelopoiesis induced by chemotherapy. We found that CTX-induced monocytes also can promote tumor metastasis in a breast cancer mouse model (Fig. 5). Although we did not conduct transfer experiment using sorted NeuMo cells due to the technical difficulty of collecting sufficient number of cells, we suspect that the NeuMo subset was the primary driver of tumor metastasis because it is more prevalent in the CTX-induced monocyte population (Fig. 3B). The immunosuppressive activity and metastasis-promoting capability of the NeuMo cells suggest the potential role of these cells in promoting tumor relapse. Our findings raise the possibility that targeting the pro-tumor NeuMo cells may improve the antitumor outcome of chemotherapy and immunotherapy.

The limitation of our study is that we only characterized chemotherapy-induced NeuMo cells in mice. Although the human counterpart of NeuMo cells have been identified in the peripheral blood of healthy donors(17), it awaits to be verified that NeuMo cells can be found in patients receiving chemotherapy and these NeuMo cells have pro-tumor activities. Cyclophosphamide, gemcitabine, paclitaxel and doxorubicin are widely used in the clinic for the treatment of various types of cancer. Therefore, it is important to investigate in cancer patients the presence, composition and function of the myeloid cells repopulating after chemotherapy. Our findings obtained in mice may inform future translational studies and lead to the development of more effective treatment strategies for patients with cancer.

## Acknowledgments

We thank Rebekah Tritz and David Hansen at the Georgia Cancer Center Flow and Mass Cytometry Core for cell sorting. We thank Miao Yu and Martina Zoccheddu at The Integrated Genomics Shared Resource in the Georgia Cancer Center for single-cell RNA sequencing. We thank the Georgia Cancer Center Summer Research Experience (SRE) Program for sponsoring this study.

## Author Contributions

Z-C.D. performed research and analyzed data; G.I.Z. conducted scRNAseq bioinformatics analysis; O.D.O, and X.W. assisted experiments; L.B. advised study design and provided clinical samples; G.Z. provided funding, designed research and wrote the paper. H.S. provided funding, designed research, analyzed data and wrote the paper.

